# Metagenomic analysis of coprolites from three Late Pleistocene megaherbivores from the Southwestern United States

**DOI:** 10.1101/2022.05.06.490351

**Authors:** Tomos O. Prys-Jones, Tara N. Furstenau, Andrew J. Abraham, Isaac N. Shaffer, Colin J. Sobek, Jordyn R. Upton, Samantha N. Hershauer, Kelvin Wong, Marirosa Molina, Sebastian Menke, Jim I. Mead, Christopher H. Ebert, Mariah S. Carbone, Edward A.G. Schuur, Faith M. Walker, Viachelsav Y. Fofanov, Christopher E. Doughty

**Affiliations:** School of Informatics, Computing and Cyber Systems, Northern Arizona University, Flagstaff, Arizona, USA; Pathogen and Microbiome Institute, Northern Arizona University, Flagstaff, Arizona, USA; Office of Research and Development, US Environmental Protection Agency, Durham, North Carolina, USA; Evolutionary Ecology and Conservation Genomics, University of Ulm, Ulm, Germany; The Mammoth Site, Hot Springs, South Dakota, USA; Desert Laboratory at Tumamoc Hill, University of Arizona, Tucson, Arizona, USA; Center for Ecosystem Science and Society, Northern Arizona University, Flagstaff, Arizona, USA

**Keywords:** megafauna, mammoth, bison, ground sloth, metagenomics, microbiome, coprolite, desiccation

## Abstract

1.

**Background:** Determining the life-history traits of extinct species is often difficult from skeletal remains alone, limiting the accuracy of studies modeling past ecosystems. However, the analysis of the degraded endogenous bacterial DNA present in paleontological fecal matter (coprolites) may enable the characterization of specific traits such as the host’s digestive physiology and diet. An issue when evaluating the microbial composition of coprolites is the degree to which the microbiome is representative of the host’s original gut community versus the changes that occur in the weeks following deposition due to desiccation. Analyses of paleontological microorganisms are also relevant in the light of recent studies linking the Late Pleistocene and Early Holocene extinctions with modern-day zoonotic pathogen outbreaks.

**Methods:** Shotgun sequencing was performed on ancient DNA (aDNA) extracted from coprolites of the Columbian mammoth (*Mammuthus Columbi*), Shasta ground sloth (*Nothrotheriops shastensis*) and paleontological bison (*Bison sp.*) collected from caves on the Colorado Plateau, Southwestern USA. The novel metagenomic classifier MTSv, parameterized for studies of aDNA, was used to assign bacterial taxa to sequencing reads. The resulting bacterial community of coprolites was then compared to those from modern fecal specimens of the African savannah elephant (*Loxodonta africana*), the brown-throated sloth (*Bradypus variegatus*) and the modern bison (*Bison bison*). Both paleontological and modern bison fecal bacterial communities were also compared to those of progressively dried cattle feces to determine whether endogenous DNA from coprolites had a microbiome signal skewed towards aerobic microorganisms typical of desiccated fecal matter.

**Results:** The diversity of phyla identified from coprolites was lower than modern specimens. The relative abundance of Actinobacteria was increased in coprolites compared to modern specimens, with fewer Bacteroidetes and Euryarchaeota. Firmicutes had a reduced relative abundance in the mammoth and bison coprolites, compared to the African savanna elephants and modern bison. There was a significant separation of samples in NMDS plots based on their classification as either paleontological or modern, and to a lesser extent, based on the host species. Increasingly dried cattle feces formed a continuum between the modern and paleontological bison samples.

**Conclusion:** Our results reveal that any coprolite metagenomes should always be compared to desiccated modern fecal samples from closely related hosts fed a comparable diet to determine the degree to which the coprolite metagenome is a result of desiccation versus true dissimilarities between the modern and paleontological hosts. Also, a large-scale desiccation study including a variety of modern species may shed light on life-history traits of extinct species without close extant relatives, by establishing the proximity of coprolite metagenomes with those from dried modern samples.

## 2. INTRODUCTION

Terrestrial ecosystems have lost a significant proportion of their megafauna (species > 44.5 kg) (Stuart 2015; Malhi et al. 2016; Smith et al. 2016; Smith and Lyons 2011; Smith et al. 2018), with many of these extinctions occurring during the Late Pleistocene due to anthropogenic pressures (Smith et al. 2019; Sandom et al. 2014; Barnosky et al. 2004) and climate change (Barnosky et al. 2004; Mann et al. 2019; Monteath et al. 2021; Wang, Zhang, and Kong 2021). Studies have shown that such dramatic extinctions of megafauna have had long-lasting effects on forest structure (Malhi et al. 2016; Doughty, Faurby, and Svenning 2016), plant geographic ranges (Doughty et al. 2016; Janzen and Martin 1982; Blake et al. 2009; Beaune et al. 2013), carbon storage (Doughty et al. 2016), maintenance of high latitude grasslands (Murchie et al. 2021; Zimov et al. 2015; Bakker et al. 2016), nutrient cycling (Doughty 2017; Wolf, Doughty, and Malhi 2013; Doughty et al. 2016; Doughty, Wolf, and Malhi 2013), the perceived host specificity of parasites (Farrell et al. 2021) and emergent zoonotic diseases (Doughty et al. 2020). Additionally, the life-history traits of the megafauna are not yet classified, limiting our understanding of these big species and the accuracy of studies modelling their impact.

It is possible to detect life-history traits in extant hosts through an analysis of the microbiome, the community of microbes present on and within an organism that is known to be important in many aspects of host health (Mansour et al. 2021; Grice and Segre 2011) and behavior (Ezenwa 2003; Johnson and Foster 2018). The microbiome composition varies with host phylogeny (Kartzinel et al. 2019), intra-species genetic variation (Blekhman et al. 2015; Goodrich et al. 2014), sex (Blyton et al. 2014), host age (Gordon, Stern, and Collignon 2005), size (Gordon and Cowling 2003), diet (Groussin et al. 2017; Carmody et al. 2015; Kartzinel et al. 2019), gut physiology (Ley et al. 2008), geography (Hartel et al. 2002), other microbiome species (Gordon, O’Brien, and Pavli 2015), transmission of microbes from other hosts (VanderWaal et al. 2014; Blyton et al. 2014) and external sources (Chiyo et al. 2014), and due to anthropogenic factors such as antibiotic usage (Langdon, Crook, and Dantas 2016). Determining such life-history traits from extinct species may also be possible through an analysis of paleontological microbiomes and may resolve on-going debates such as the gut physiology of the extinct ground sloths, which made up a significant portion of the lost Late Pleistocene megafauna (Stuart 2015). While the remaining smaller, arboreal members of the sloth Folivora sub-order (*Bradypus* and *Choloepus* sp.) have a simple foregut fermenting GI tract, metabolic scaling studies suggest that ground sloths may have employed a different digestive strategy to reach their large size (Tejada et al. 2021; Clauss et al. 2003). Abraham et al. (2021), have shown that machine learning methods using host traits, including digestive physiology, improve estimates of gut passage time, meaning that past estimates of nutrient dispersal would be improved with information on the digestive physiology of extinct species, compared to the current estimates based on allometric scaling (Wolf, Doughty, and Malhi 2013).

Characterizing paleontological microorganisms is also relevant in light of recent studies linking Late Pleistocene and early Holocene extinctions with modern-day parasites (both single- and multi-cellular) (Doughty et al. 2020; Farrell et al. 2021). In particular, Doughty et al. (2020) showed that the extinction of large mammals was associated with an increased risk of zoonotic emergent infectious diseases (EIDs) in contemporary human populations (Doughty et al. 2020). The authors hypothesized that the loss of the biggest, furthest ranging species reduced the dispersal ability of microbes (via direct transmission or ectoparasites), potentially leading to greater microbe heterogeneity across the landscape due to divergent evolution, as well as greater immune-naiveté in the remaining hosts. With the declining megafauna populations, they also suggest a selective pressure for the parasites to find new hosts, a hypothesis based on published host-parasite theory (Hoberg and Brooks 2008). Further work is needed to test these hypotheses and determine how they might link to present-day EIDs. A first step in understanding whether Late Pleistocene extinctions perturbed microbe dynamics is through a comparison of modern and paleontological microbe communities. However, there is currently a lack of studies on Late Pleistocene gut microbiomes using the latest metagenomic techniques.

Recent advances in sequencing technology and bioinformatic methods enable the investigation of ancient genomes (Poinar et al. 2006; Delsuc et al. 2019; Karpinski, Mead, and Poinar 2017). Earlier studies used polymerase chain reaction (PCR) amplification to assess a limited number of species, but the advent of shotgun metagenomics has allowed for the characterization and analysis of whole communities (Larsen, Cole, and Worobey 2018). The quality of such studies depends largely on the type of sample (e.g. bone, tooth, coprolites, masticated birch pitch) (Kashuba et al. 2018; Jensen et al. 2019; Hansen et al. 2017), as well as the conditions in which they were discovered and subsequently stored, as both these factors affect the accumulation rate of age-related damage in the DNA (Dabney, Meyer, and Pääbo 2013).

While there are many microbiome analyses in the contexts of human health (Velloza and Heffron 2017; Buvé et al. 2014; Ursell et al. 2012; Mansour et al. 2021; Banerjee et al. 2022; Grice and Segre 2011), with a growing literature in non-human animals (Ilmberger et al. 2014), studies of Late Pleistocene microbiomes are limited. Initially, these began with amplicon-based methods (Tito et al. 2012; Hofreiter et al. 2000) and permafrost specimens (Hagelberg, Hofreiter, and Keyser 2015; Mardanov et al. 2012; Ravin, Prokhortchouk, and Skryabin 2015). More recently, shotgun metagenomics has been used to analyze all the DNA present within ancient samples without relying on PCR amplification, avoiding the associated biases, and giving a more accurate insight into the relative abundances of community members (McLaren, Willis, and Callahan 2019; Ferrari et al. 2018; Durazzi et al. 2021). Although shotgun metagenomics has been performed on ancient packrat (*Neotoma* sp.) middens from the southwestern USA (Moore et al. 2020) and on Holocene coprolites (Wibowo et al. 2021; Witt et al. 2021; Rampelli, Turroni, Debandi, et al. 2021b), to the best of the authors’ knowledge, no such techniques have been applied to characterize the microbiomes of desiccated megafauna coprolites from the Late Pleistocene (epoch end date: 11,700 ybp) (Gradstein, Ogg, and Smith 2005). Although not from coprolites directly, a notable study by Rampelli et al (2021) identified commensal gut microbes from sediments at a Spanish site occupied by Neanderthals around 50 kybp (Rampelli, Turroni, Mallol, et al. 2021). The sediments contained fecal lipid biomarkers.

Coprolites are readily preserved on the Colorado Plateau, in the U.S. Southwest, due to the aridity of the environment (Reheis et al. 2005) and the abundance of cave systems that provide shelter from precipitation (Hansen 1978; Mead and Swift 2012). The Plateau measures roughly 337,000 km^2^, has an average elevation of 1525 m and spans parts of Arizona, Colorado, New Mexico, and Utah (Anderson et al. 2000; Martin, Martin, and Mead 2017). Walker et al. (2019) recorded a stable, low humidity and temperature at a cave site near Glen Canyon, Utah, over a three-year period, both favorable conditions for DNA preservation. The desiccated fecal matter found in the caves originates from multiple megafauna species (Mead and Agenbroad 1992), with the most common contributors being Columbian mammoth (*Mammuthus columbi*) (Mead and Agenbroad 1992; Mead and Agenboard 1989; Mead et al. 1986), Shasta ground sloth (*Nothrotheriops Shastensis*) (Mead and Agenbroad 1992; Mead and Agenboard 1989; Long, Hansen, and Martin 1974; Schmidt, Duszynski, and Martin 1992), bison species (*Bison sp.*) (Mead and Agenboard 1989; Mead and Agenbroad 1992), camels (*Camelops* sp.) (Mead and Agenbroad 1992), native horses (*Equus* sp.) (Mead and Agenboard 1989), bighorn sheep (*Ovis canadensis*) (Mead and Agenbroad 1992; Campos et al. 2010), and Harrington’s mountain goat (*Oreamnos harringtoni*) (Mead and Agenboard 1989; Mead and Agenbroad 1992; Mead, O’Rourke, and Foppe 1986; Mead et al. 1987).

In many cases, intact boluses from multiple species were preserved in extensive blankets of feces, together with hair from the same species. At Bechan Cave, Utah, the blanket measured 255 m^2^ and was estimated to contain 370 m^3^ of organic material (Davis et al. 1984). Alongside the coprolites are often preserved packrat middens (Mead, Thompson, and Long 1978; Moore et al. 2020; Anderson et al. 2000) – nest structures created by the extant packrat, in which animal and plant material are held together in a cement of crystalized urine and that are recognized as preserving genetic material (Wilder et al. 2014; Mead et al. 2021; Davis et al. 1984a).

Since the early twentieth century, the macrobotanical, microhistological and pollen samples found within these coprolites and middens have been used to reconstruct the environment of the Southwest USA, and in the case of the fecal matter, the diet of the originator (Eames 1930; Martin, Sabels, and Shutler 1961; Hansen 1978; Thompson et al. 1980; Martin, Thompson, and Long 1985; Mead, O’Rourke, and Foppe 1986; Mead et al. 1987; Schmidt, Duszynski, and Martin 1992). Later genetic analyses of the Colorado Plateau coprolite specimens have focused on amplifying and sequencing gene fragments from the host and dietary plants it consumed (Hofreiter et al. 2000; Poinar et al. 1998; Greenwood et al. 2001; Thompson et al. 1980; Mead, Schroeder, and Yost 2021). Poinar et al. (1998) identified boluses found in Gypsum Cave, Nevada, as originating from the *Xenarthra* taxon and subsequently assigned them to the Shasta ground sloth by sequencing multiple fragments of mitochondrial rDNA genes, as well as amplifying a fragment of the rbcL gene to identify the dietary plant families (Poinar et al. 1998; Hofreiter et al. 2000). Similarly, Karpinski, Mead, and Poinar (2017) were able to confirm that the boluses identified at Bechan Cave came from the Columbian mammoth, concurring with previous morphometric analysis done by Mead et al. (1986).

For fecal matter deposited into an arid environment, such as that found on the Colorado Plateau, there exists a window of time in which the bolus’ microbial species composition changes, such that certain species are lost, while others change their abundances in a predictable direction (Menke, Meier, and Sommer 2015; Wong et al. 2016; Oladeinde et al. 2014). Desiccation eventually limits the risk of external contamination, as the bacterial growth is reduced when the water content falls (Sinton et al. 2007; Merino et al. 2019) and even stops as the relative humidity of the air surrounding the growth medium drops below 60% (Beuchat et al. 2013; Esbelin, Santos, and Hébraud 2018). However, during the time when enough moisture remains, exposure to oxygen and UV radiation changes the microbial composition of the specimen (Menke, Meier, and Sommer 2015). The obligate anaerobes that grow in the anoxic conditions of the gut are often lost or reduced in abundance as fecal matter is exposed to atmospheric oxygen concentrations. Therefore, coprolites containing endogenous DNA are likely to have a microbiome signal skewed towards aerobic microorganisms typical of desiccated fecal matter, rather than being representative of the original gut community.

Here we classified the commensal organisms present within the gastrointestinal (GI) tracts of three Late Pleistocene megafauna species native to the United States Southwest through an analysis of coprolites using metagenomic shotgun sequencing and a novel metagenome analysis pipeline. These samples were collected on the Colorado Plateau and include the now extinct Columbian mammoth, Shasta ground sloth, as well as North American bison species. The main objectives of this study are a) to characterize the organisms present within coprolites of the three Late Pleistocene species, b) to compare these communities to those found in the feces of closely related modern day species where available, and c) to assess whether the coprolites have a microbiome signal typical of desiccated fecal matter.

## 3. MATERIALS AND METHODS

### 3.1 FIELD RESEARCH PERMITS AND PERMISSIONS

All samples were collected with the approval of the appropriate National Park Services. Mammoth, Shasta ground sloth and paleontological bison coprolites were available under permits: #L.2017.27, #GRCA-2018-SCI-0030, and #GLCA-2019-SCI-0010, respectively, made to the Glen Canyon and Grand Canyon National Parks. Modern bison fecal samples were collected under permit (#GRTE-2019-SCI-0058, made with the Grand Teton National Park, and the elephant boluses were collected with the approval of Zambia’s Department of National Parks and Wildlife (permit #0043037).

### 3.2 COPROLITE AND MODERN FECAL SAMPLES

Paleofecal matter from the Shasta ground sloth, Columbian mammoth and bison were obtained from the Museum of Northern Arizona (Flagstaff, USA) and the Grand Canyon National Park Museum (S1 Table). The five fragments of paleofeces from bison were found in three locations in Glen Canyon National Park Recreation Area, Utah: Mammoth Alcove (three fragments; n = 3), Grobot Grotto (n = 1) and Hooper’s Hollow (n = 1) (Mead and Agenbroad 1992; Mead and Agenboard 1989; Martin, Martin, and Mead 2017). The six mammoth coprolites included here were collected from Bechan cave, Utah (Jim I. Mead et al. 1986; J. Mead and Agenboard 1989; Paul S. Martin 1987; Agenbroad and Mead 1989), and the Shasta ground sloth samples came from two locations in Arizona, Muav (n = 3) and Rampart Caves (n = 3) (Schmidt, Duszynski, and Martin 1992; Hofreiter et al. 2000; Poinar et al. 2003; Poinar et al. 1998; Long and Martin 1974; Martin, Sabels, and Shutler 1961; Hansen 1978; Martin, Thompson, and Long 1985) (example of bolus in Figure 1).

**Figure 1.**
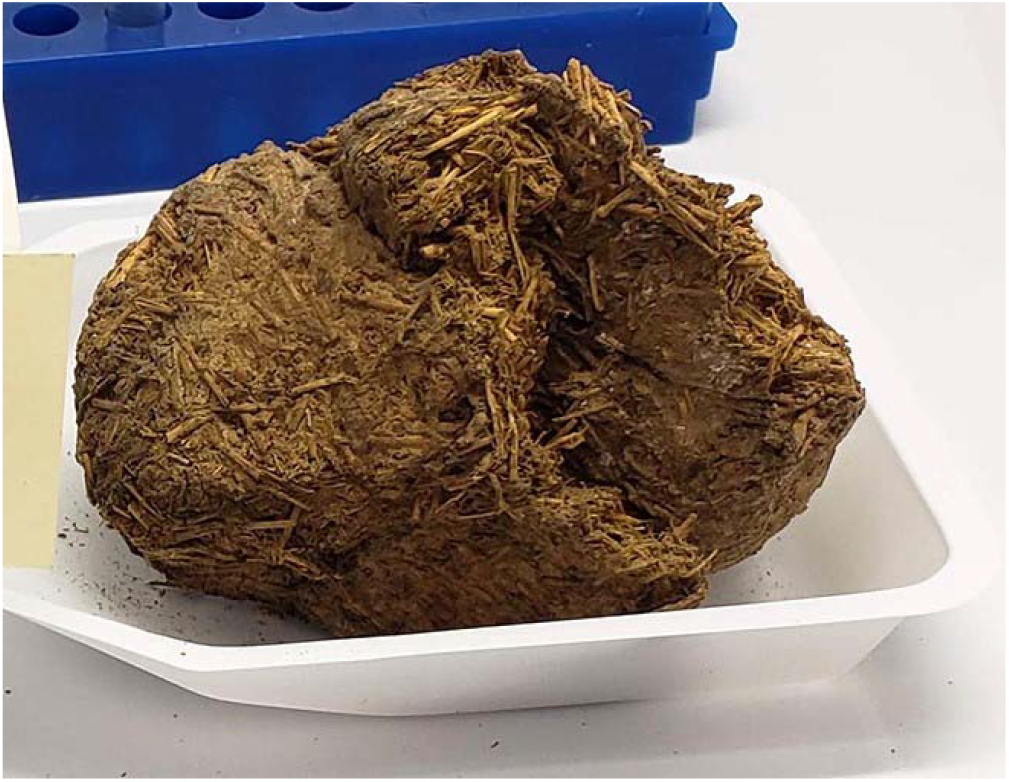
Coprolite bolus from the Shasta ground sloth (*Nothrotheriops shastensis*) collected from Rampart Cave. Further sample details provided in Table S1.

The samples were prepared for 14C analysis in 2021 at the Arizona Climate and Ecosystem (ACE) Isotope Laboratory at Northern Arizona University. The interior of each blouse was subsampled and homogenized using a mortar and pestle, and approximately 2.5 mg of dry organic matter was weighed into a tin capsule and converted to graphite using the Automated Graphitization Equipment (AGE 3, Ionplus, Switzerland). The 14C content of the graphite was measured using accelerator mass spectrometry (AMS) on a Mini Carbon Dating System (MICADAS, Ionplus, Switzerland). The data are reported in fraction modern following standard methods in Trumbore, Sierra, and Hicks Pries (2016). Instrument error is reported for all 14C data; and ranged between 0.32 – 0.43% for all samples. The radiocarbon ages were converted to calendar date using the OxCAL model version 4.4 (Ramsey 2009) and atmospheric data from Reimer et al. (2020). A single replicate was radiocarbon dated for each of the mammoth and bison coprolites, whereas multiple replicates were taken from the sloth boluses.

Modern fecal matter was also investigated with the same pipeline used to categorize the metagenomes of the coprolites, to compare the microbes present in samples from related hosts. These samples were selected from related species that had a comparable diet and, where possible, gut type. The natural comparison to coprolites from paleontological bison were fresh samples collected in October 2019 from five free-ranging bison (*Bison bison*) in the Grand Teton National Park, Wyoming. Samples were taken as soon as was safely possible (within < 2 hrs. post deposition), moved to a portable freezer in the field and then transported to the laboratory on dry ice. Counterparts to mammoth coprolites were two specimens of dung from African savannah elephants (*Loxodonta africana*) collected in June 2018 from wild individuals in the Nsefu sector of South Luangwa National Park, Zambia, as well as four fecal metagenomes downloaded from NCBI (ERR4083547, ERR4083568, ERR4083765 and ERR4084019). A portion of each Zambian bolus was placed into a 15 mL conical tubes, covered with RNALater solution, and then frozen -20 °C before being moved to the laboratory for further processing. The elephants were not observed defecating, as with the bison samples, but were verified as the originator by amplifying the cytochrome oxidase subunit I (COI) with VF1/VR1 primers using recommended PCR conditions (Ivanova, Clare, and Borisenko 2012).

All surviving species within the *Folivora* sub-order of sloths have distinct life-history traits to the ground sloths (Urbani and Bosque 2007; Hayssen 2010). Although no fresh fecal samples were processed from extant members of the sloth *Folivora* sub-order, two fecal metagenomes from the brown-throated sloth (*Bradypus variegatus)* were downloaded from NCBI (ERR4083577 and ERR4083789) – the closest extant sloth genus to the Shasta ground sloth (Delsuc et al. 2019).

### 3.3 DNA EXTRACTION, SEQUENCING, AND ADAPTER REMOVAL

The extraction of genetic material from coprolites was performed at a laboratory dedicated to processing ancient DNA, based at the Northern Arizona University. Material from the center of each coprolite bolus was taken in order to minimize the risk of contamination from external sources (Wood and Wilmshurst 2016), and was extracted using a modified phenol-chloroform protocol described by Fleischer et al. (2000). The Fleischer et al (2000) method was adjusted during DNA extraction such that the Amicon elution step consisted of the specified volume of water being spun down over the course of two cycles of 30 minutes using a Fisherbrand 614B Series Centrifuge, resulting in a final elution volume measuring approximately 150 µL. For modern samples, approximately 0.25 mg of feces was added into 1.5 mL microcentrifuge tubes, subjected to a quick spin, before the RNAlater was pipetted off. A QIAamp Fast DNA Stool Mini Kit (Qiagen, Hilden, Germany) was then used according to the manufacturer protocol for human analysis (‘*Isolation of DNA from Stool for Human DNA Analysis*’).

DNA library construction for whole-genome sequencing was performed using the KAPA Hyper Prep Kits for Illumina® NGS platforms (KAPA Biosystems, KK8504). Adapters and 8bp index oligos purchased from IDT® (Integrated DNA Technologies, San Diego, CA) based on Kozarewa and Turner (2011), were used in place of those supplied in the KAPA preparation kit (Kozarewa and Turner 2011). Modifications were made to the post-adapter ligation clean-up, indexing PCR and post-PCR clean-up to account for the low input DNA concentrations. The final libraries were quantified on an Applied Biosystems™ QuantStudio™ 7 Flex Real-Time PCR System using the KAPA SYBR® FAST ROX Low qPCR Master Mix for Illumina platforms (KAPA Biosystems, KK4873). DNA extracted from mammoth and sloth boluses were sequenced on an Illumina MiSeq using the 600-cycle v3 kit (Illumina, MS-102-3003), whereas the elephant and modern and paleontological bison samples were sequenced on an Illumina NextSeq 500 system. Both platforms were run using standard Illumina procedures and the appropriate sequencing primers were added to the kits as in Kozarewa and Turner (2011). FastQC version 0.11.9 was used to assess the quality of the generated readsets and, when detected (Andrews 2010), adapter contamination was removed using Cutadapt version 3.2 (M. Martin 2011).

### 3.4 TAXONOMIC CLASSIFICATION

Metagenomic sequencing reads were categorized using a novel metagenomic classifier (MTSv version 2.0.0) (Fofanov, Viacheslav Y.Furstenau et al. 2017) run using the Monsoon high performance computing cluster based at Northern Arizona University. MTSv is a taxonomic assignment tool that performs a full alignment with reference sequences using an FM-index assisted q-gram filter (to quickly find likely alignment locations) followed by SIMD accelerated Smith-Waterman alignment (Ferragina and Manzini 2000; Reinert et al. 2015; Rasmussen, Stoye, and Myers 2006; Burkhardt et al. 1999; Zhao et al. 2013). The FM-indices were built using reference sequences downloaded from the complete GenBank repository on October 28, 2019. Reads that passed the default quality control parameters of the Fastp toolkit (Chen et al. 2018) were then partitioned into shorter k-mer queries and then deduplicated (whilst retaining the number of query copies) to reduce redundant alignments. Overlapping substrings of the same size (seed size) were extracted from each query (and its reverse complement) at specified intervals (seed gap). The position of exact seed matches was looked up using the FM-index and locations that had enough seed hits (specified by the min-seed cut-off) became candidates for full alignment. If the edit distance of the full alignment was less than or equal to the edit distance cut-off, the query was assigned to the corresponding taxa.

When analyzing the taxonomic assignments, we identified queries that were assigned to a single taxon in the reference database and refer to these as *unique signature hits (USHs)*. USH provide the strongest support that the taxon was present in the sample. To provide further support that the reads originated from the reference taxon (e.g., rather than a related taxon that is not in the reference database) we compared the difference between expected and observed values for the ratio of USH to total unique hits. The expected values were calculated by randomly sampling 100,000 k-mers from the sequence database for a candidate taxon. The queries were then run through the same process, using the same parameters as the observed sample data, to calculate the ratio. We used the two one-sided tests (TOST) procedure to test for equivalence within an upper and lower equivalence bound that is specified based on the user-specified effect size (Cohen’s *h,* default=0.5). When both one-sided tests were rejected (α=0.05), we concluded that the difference between the expected and observed values were close enough to be practically equivalent (Schuirmann 1987; Cohen 2013).

### 3.5 MTSV ALIGNMENT PARAMETER SELECTION FOR ANCIENT DNA

The read sequences from organisms present in coprolites have the potential to differ significantly from the genomes of predominantly modern organisms in the reference database, due to a combination of divergent evolution and age-related degradation of the genetic material. To test the most favorable parameters to classify aDNA using MTSv, we introduced age-related changes into 1,000,000 reads generated in-silico from 200 randomly selected firmicute species using the gargammel simulator (Renaud et al. 2017), prior to classification. Firmicutes were selected as an appropriate bacterial phylum as they are often a major component of herbivore fecal microbiomes (Donnell et al. 2017). To account for the variation in bacterial abundances in herbivore GI tracts, the proportion of reads assigned to each bacterial species was random and ranged linearly between 173 and 24,383. Such low read numbers were used to represent the most extreme edge cases of degradation in paleontological samples.

Gargammel generates reads in prespecified proportions from selected FASTA files then cuts and changes the base composition of the reads to represent the distribution of read lengths and deamination patterns appropriate for paleontological samples, respectively. Adapters are then added to the raw Illumina reads and sent to ART NGS simulator to include sequencing errors and corresponding quality scores (Huang et al. 2012). In this case reads lengths were distributed according to Fu et al. (2014) and deaminated according to the Briggs et al. (2007) model. Data from Fu et al (2014) and Briggs et al (2007) were used owing to their independence from the present study and being verified paleontological sources of DNA with minimal contamination.

The accuracy of MTSv in correctly classifying the gargammel reads to the class, order, family, genera, and species levels was then assessed using the positive predictive value (PPE) and sensitivity score (PPE = true positives/ true positives + false positives; sensitivity = true positives / true positives + false negatives). The pipeline was run with different combinations of parameters, varying the query k-mer size, the number of permitted edits between the query and reference, as well as the sensitivity of the seed filter (parameter combinations summarized in S2 Table). MTSv outputs all taxa detected with at least a single USH, leading to many taxa being identified with very few hits. In such cases there is uncertainty on whether the identified taxa were present. As a result, a minimum USH threshold was also tested for the inclusion of taxa, with the threshold ranging from 0 to 1800 USH (in increments of 200 USH). This was then converted into a proportion of the total USH.

### 3.6 MTSV OUTPUT AND BETA DIVERSITY METRICS

Many studies comparing community structure between different samples rely on beta diversity metrics, accounting for differences in species richness and often abundance, and which allow for subsequent ordination analyses. To determine whether these metrics could be used with the MTSv output, the relative abundance of reads from taxa input into gargammel was compared to the proportion of USH that MTSv assigned to the same taxa. This analysis was repeated at the class, order, family, genus, and species levels for the 200 firmicutes. Two metrics were used to assess the correlation, Lin’s concordance correlation coefficient (ρ_c_) (Lin 1989) and the R^2^ value of points around a 1-to-1 line. Only taxonomic levels that had a tight correlation between gargammel read proportions and the relative abundance of USH from MTSv, satisfying McBride’s interpretation of an almost perfect correlation (ρ_c_ > 0.99) (Akoglu 2018; McBride et al. 2005), were used in the subsequent comparison of the coprolite and modern samples. Bray-Curtis dissimilarities were calculated from the bacterial communities in samples using the vegan package version 2.5-6 in R 4.0.3 (Oksanen et al. 2019). This metric was selected due to its relative insensitivity to undersampling and taxonomic errors (Schroeder and Jenkins 2018).

### 3.7 DEAMINATION

Following the completion of the MTSv runs, age-related DNA damage was analyzed in the reads associated with the identified taxa, to assess whether significant modern contamination was introduced into the coprolite samples during preparation. Modern contamination of ancient samples is typified by higher concentrations of DNA than the background of highly degraded endogenous genetic material (low coverage and depth). Ancient DNA is also highly fragmented, with a characteristic pattern of nucleotide modifications (Pääbo 1989; Dabney, Meyer, and Pääbo 2013; Skoglund et al. 2014), even in frozen (Ravin, Prokhortchouk, and Skryabin 2015; Mardanov et al. 2012; Ferrari et al. 2018; Van Geel et al. 2011) or fully desiccated (Hofreiter et al. 2000; Karpinski, Mead, and Poinar 2017; Poinar et al. 2003; Delsuc et al. 2019; Wood et al. 2013) specimens that tend to encourage DNA preservation.

Queries associated with each species identified by MTSv, which had over the threshold proportion of USH, were separated using the pipeline’s extract functionality. Any NCBI taxonomic identification numbers for ‘*Homo sapiens*’, ‘uncultured bacterium’ and ‘synthetic construct’ were excluded. The original reads were then identified from these queries and subsequently aligned against the reference genomes where available, using the Burrows-Wheeler Algorithm (bwa aln), parameterized to optimize the alignment of short read aDNA (- l 1024 -n 0.01 -o 2) (Oliva et al. 2021). The output SAM files from the alignment were then processed using MapDamage2 (Jónsson et al. 2013), a tool to assess the age-related C-to-T and A-to-G base pair transition at read 5’ and 3’ termini, respectively. Owing to the low number of reads per taxon, and therefore infrequency of detected nucleotide changes in the terminal 25 bp per read, the records of C-to-T and A-to-G transitions were summed across taxa. This was done by weighting the score for each taxon by the relative proportion of reads from that taxon, to increase the influence of any potential contamination (expected to be at higher concentrations), and therefore its detectability if present.

### 3.8 DESICCATION DATA

To determine whether endogenous DNA present in the Late Pleistocene bison coprolites had a microbiome composition more similar with dried modern fecal matter, data were included from two desiccation studies by Wong et al. (2016) and Menke, Meier, and Sommer (2015). Both assessed how the microbial composition of fresh ungulate fecal samples changed with exposure to conditions outside of the GI tract. It was then possible to determine whether the fresh bison samples were more like fresh ungulate feces regarding microbiome composition, and paleontological bison samples closer to desiccated ungulate feces. Wong et al. (2016) sampled shaded and unshaded cattle feces on days 0, 2, 4, 6, 8, 15, 22, 29, 43 and 57, amplified the 16S rRNA hypervariable V4 region and classified the reads using QIIME version 1.8.0. Operational Taxonomic units were selected using UCLUST with a 3% threshold and classified using the RDP 2.2. Menke et al (2015) used a similar methodology, sampling two springbok (*Antidorcas marsupialis*) and two giraffe (*Giraffa camelopardalis*) fecal samples daily on days 1-7, amplifying the same region of 16S rRNA and using the QIIME classifier. For both studies, the relative abundances of identified taxa were grouped with those from the paleontological and bison samples and the Bray-Curtis dissimilarities calculated. The species included in the studies each represent a different foraging strategy on the grazer (cattle) – mixed feeder (springbok) – browser (giraffe) spectrum.

### 3.9 ORDINATION AND STATISTICS

Nonmetric multidimensional scaling (NMDS) was performed based on the Bray-Curtis dissimilarities between paleontological and modern samples, as well as between the modern bison samples, paleontological bison coprolites and desiccated samples from either Wong et al. (2016) or Menke, Meier, and Sommer (2015) using the vegan package version 2.5-6 (Oksanen et al. 2019) in R 4.0.3. The same package was used to perform a permutational multivariate analysis of variance (PERMANOVA), assessing the contributions of age and host species in describing the dissimilarity between samples, with the host species nested within the host order when multiple orders were compared. Pairwise PERMANOVA were also performed to determine which age categories of cattle feces differed significantly from the paleontological versus modern bison boluses, using the Benjamini and Yekutieli (2001) p-value correction method for multiple comparisons.

## 4. RESULTS

### 4.1 COPROLITE ^14^C-DATES

Radiocarbon dates established that the bison and mammoth coprolites were older than 11,700 ybp, recognized as the end of the Late Pleistocene, whereas coprolites from the Shasta ground sloth were deposited during the Early Holocene (< 11,700 ybp). The dates also separate the two samples from the neighboring Grobot’s Grotto (OxCAL 95% probability date ranges: 20,982 – 20,592 ybp) and Hooper’s Hollow (16,903 – 16,772 ybp), indicating that the coprolites came from at least two separate bison individuals (Table S1). The same was not true for the six Columbian mammoth samples from Bechan Cave, with each coprolite age range overlapping with at least one other (maximum to minimum values for OxCAL 95% probability date ranges across all coprolites: 12,992 – 12,143 ybp). The proximity of Rampart and Muav Caves (Supplementary Material) and the overlapping date ranges from the three sloth coprolites (Table S1), also prevents distinction between originators of the sloth coprolites. Interestingly, replicate measurements from the sloth boluses were non-overlapping in several cases, most likely due to noticeable heterogeneity in the sloth boluses when prepping for 14C.

### 4.2 SELECTION OF MTSV PARAMETERS & TAXONOMIC LEVELS

To test the most favorable parameters to classify aDNA using MTSv, age-related changes were introduced into reads generated in-silico from 200 randomly selected firmicute species using the gargammel simulator, prior to classification. The PPE and sensitivity scores for MTSv in classifying the gargammel reads generally decreased with progressively lower taxonomic levels, from class to species. Each MTSv run was ranked in order of descending PPE and sensitivity. For the twenty highest ranked runs at each taxonomic level, the PPV was 100% for class and order-level classification, dropping to 81.82–88.89% for family, 93.33– 100% for genera, and 81.25-88.89% for species. The sensitivity dropped more rapidly, from 100% for class-level classification, to 55.56-66.67% for order, 31.03-62.07% for family, 8.18–12.73% for genera, and 3.50-7.50% for species. Supplementary Figure 1 shows the MTSv runs ranked in descending order of PPE and sensitivity scores at the class, order, and family taxonomic levels. The low PPE and sensitivity scores in classification of genera and species excluded these levels from further analysis. Based on the gargammel simulations the preferred MTSv parameters for the classification of ancient DNA was a k-mer query length of 36 bp, 3 edits permitted between a given read and reference, and the alignment parameters set to efficient (seed size = 14; minimum seeds = 4; seed gap = 2) (highlighted in Figure S1).

To determine whether the MTSv output could be used to calculate the Bray-Curtis dissimilarities between samples, the relative abundance of gargammel reads for each taxon was also correlated against the proportion of USH identified by MTSv (Figure S2). Only taxonomic levels above family recorded a Lin’s concordance correlation coefficient (ρ_c_) greater than 0.99, McBride’s interpretation for an almost perfect correlation (McBride et al. 2005). Therefore, Bray-Curtis dissimilarities between ancient samples, and resultant NMDS plots, were only considered accurate for phylum, class (ρ_c_ = 0.997, R^2^ = 0.993), and order levels (ρ_c_ = 0.992, R^2^ = 0.985).

### 4.3 SEQUENCING & MTSV CLASSIFICATION

The paleontological samples were classified using the preferred MTSv parameters determined using gargammel, whereas the modern samples were run using default parameters (query length: 50 bp queries, edits permitted: 3, alignment sensitivity: efficient). A total of 38,099,993 reads were generated from the coprolites and 101,874,763 reads from the modern samples. From the paleontological and modern readsets, MTSv generated a total of 25,556,061 and 23,746,478 queries, respectively. A full breakdown of reads and queries per sample is available in Table 1, including read and query number for the downloaded NCBI readsets.

**Table 1.** The number of reads after quality control for each host species and sample ID, as well as the number of queries generated by MTSv.

| Host species | Sample ID | Read number (post quality control) | Query number |
| --- | --- | --- | --- |
| *Bison bison* (modern) | NHCS-101 | 12,003,784 | 11,789,656 |
|  | NHCS-102 | 19,409,212 | 18,767,318 |
|  | NHCS-103 | 21,566,990 | 21,033,185 |
|  | NHCS-151 | 17,158,696 | 16,695,661 |
|  | NHCS-152 | 18,661,410 | 18,170,952 |
| *Bradypus variegatus* | ERR4083577 | 1,302,806 | 6,552,967 |
|  | ERR4083789 | 1,098,426 | 5,682,928 |
| *Loxodonta africana* | Zam-eleA | 6,297,172 | 5,659,763 |
|  | Zam-eleB | 13,074,666 | 6,297,059 |
|  | ERR4083547 | 1,224,644 | 6,443,389 |
|  | ERR4083568 | 617,939 | 3,251,689 |
|  | ERR4083765 | 2,620,176 | 13,483,902 |
|  | ERR4084019 | 1,260,865 | 6,651,297 |
| *Bison sp.* (paleontological) | GLCA-821A | 14,346,330 | 8,406,082 |
|  | GLCA-821B | 10,904,827 | 5,686,713 |
|  | GLCA-821C | 10,661,826 | 5,686,713 |
| *Mammuthus columbi* | GLCA-0367 | 375,073 | 1,741,386 |
|  | GLCA-2578 | 86,730 | 311,783 |
|  | GLCA-2627 | 104,288 | 343,951 |
| *Nothrotheriops shastensis* | GRCA-4712 | 153,446 | 571,429 |
|  | GRCA-59574A | 205,032 | 778,160 |
|  | GRCA-59574B | 385,020 | 1,696,570 |
|  | GRCA-59574C | 50,958 | 190,853 |

### 4.4 BACTERIAL TAXA OLD AND NEW

Generally, a lower diversity of phyla and classes were observed in the paleontological samples compared to the modern counterparts, with Figure 2 displaying the breakdown per specimen. Across all coprolites the relative abundance of Bacteroidetes and Euryarchaeota were lower, compared to the relatively higher abundance of Actinobacteria (Table S3). Firmicutes were also at a generally low relative abundance in the mammoth and bison coprolites, compared to the African savanna elephants and modern bison, respectively.

**Figure 2.**
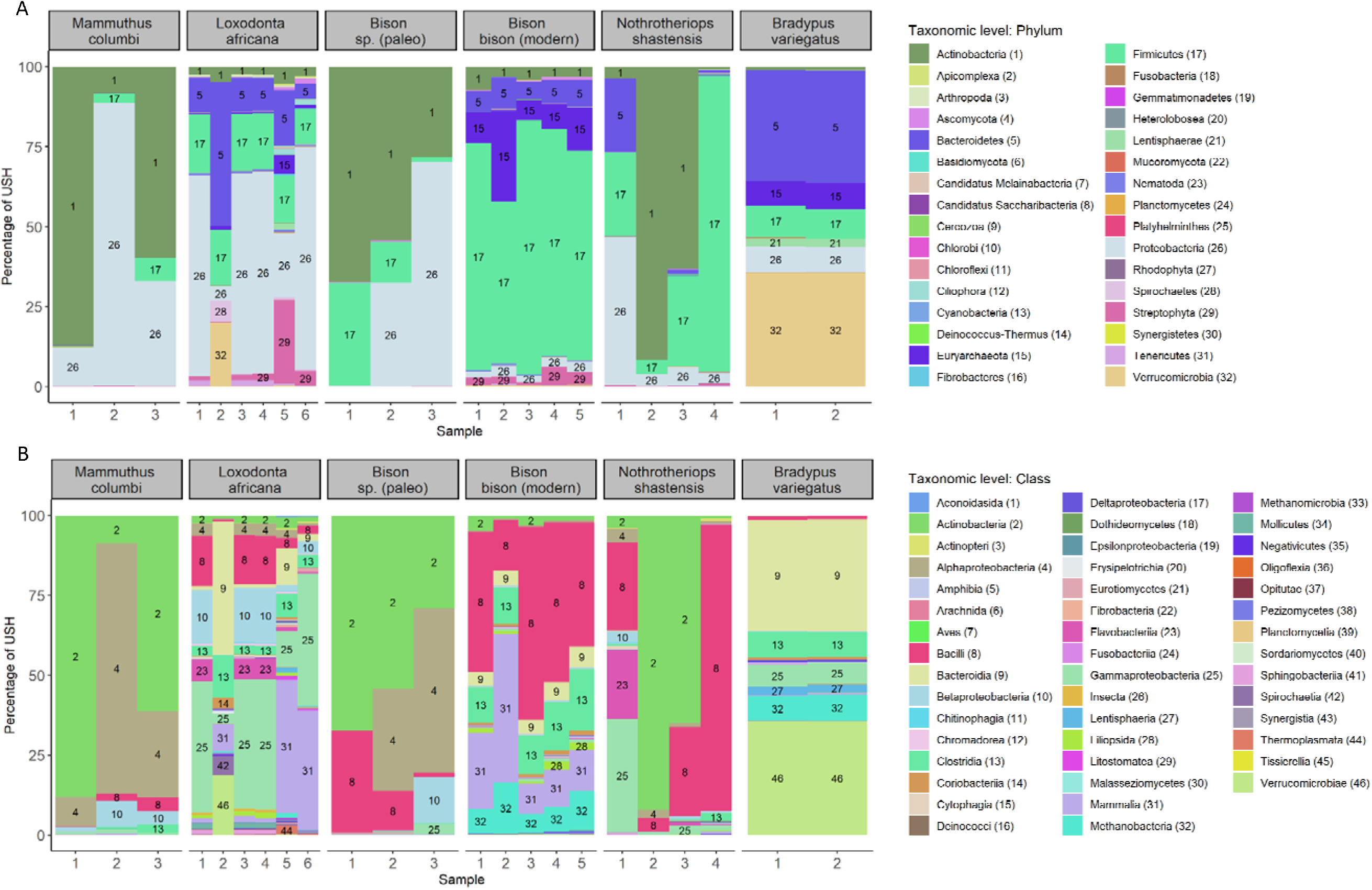
Organisms detected within coprolites from the Columbian mammoth, African savannah elephant, paleontological bison, modern bison, Shasta ground sloth and brown-throated sloth (grey boxes, left-to-right) at the phylum (A) and class (B) taxonomic levels. Numbers associated with each taxon were included to distinguish between taxa assigned similar colors. Relative abundance of each taxon is based on the percentage of unique signature hits (USH) detected by the metagenomic classifier, MTSv. Bacterial diversity at phylum and class levels is decreased in paleontological coprolites compared to modern specimens.

In both old and modern specimens, most reads mapped to bacteria, compared to other kingdoms. For phyla with over 1% of USH, Actinobacteria, Proteobacteria and Firmicutes dominated the bacterial signal from the coprolites, with bison having respective proportions of 49.9%, 34.3%, 15.5%, mammoth having 51.7%, 44.5%, 3.5% and Shasta ground sloth, 39.8%, 14.8%, 37.8% (Table S3: phyla with over 1% of USH). The ground sloth also had 6.3% of USH mapping to Bacteroidetes. These four phyla also dominated the modern samples, with the addition of Verrucomicrobia and Lentisphaerae in the brown-throated sloth, and Verrucomicrobia, Spirochaetes and Tenericutes present in the elephant samples (Figure 2 and Table S3).

For bacterial classes with over 1% USH, the core taxa across all the coprolites included Bacilli, Alphaproteobacteria, Betaproteobacteria, and Actinobacteria. Respectively, the percentage of USH in each class was 15.25%, 27.72%, 4.93%, and 50.24% for bison coprolites, 2.27%, 38.15%, 4.55%, and 52.70% for mammoth coprolites and 37.32%, 2.31%, 1.19%, and 40.38% for the Shasta ground sloth. Additionally, Gammaproteobacteria and Flavobacteria were detected in the ground sloth samples (10.14% and 5.76% of USH), with Gammaproteobacteria also detected in bison coprolites (1.22%), and Clostridia in mammoth boluses (1.13%).

At phylum and class levels NMDS ordination of the paleontological coprolites and modern specimens showed separation of samples based on their paleontological versus modern classification and the identity of the originator (Figure 3). PERMANOVA confirmed that host species (nested within host order) and age (modern versus paleontological) were significant variables in determining the dissimilarity of samples (p-value ≤ 0.001 for both variables at phylum and class levels).

**Figure 3.**
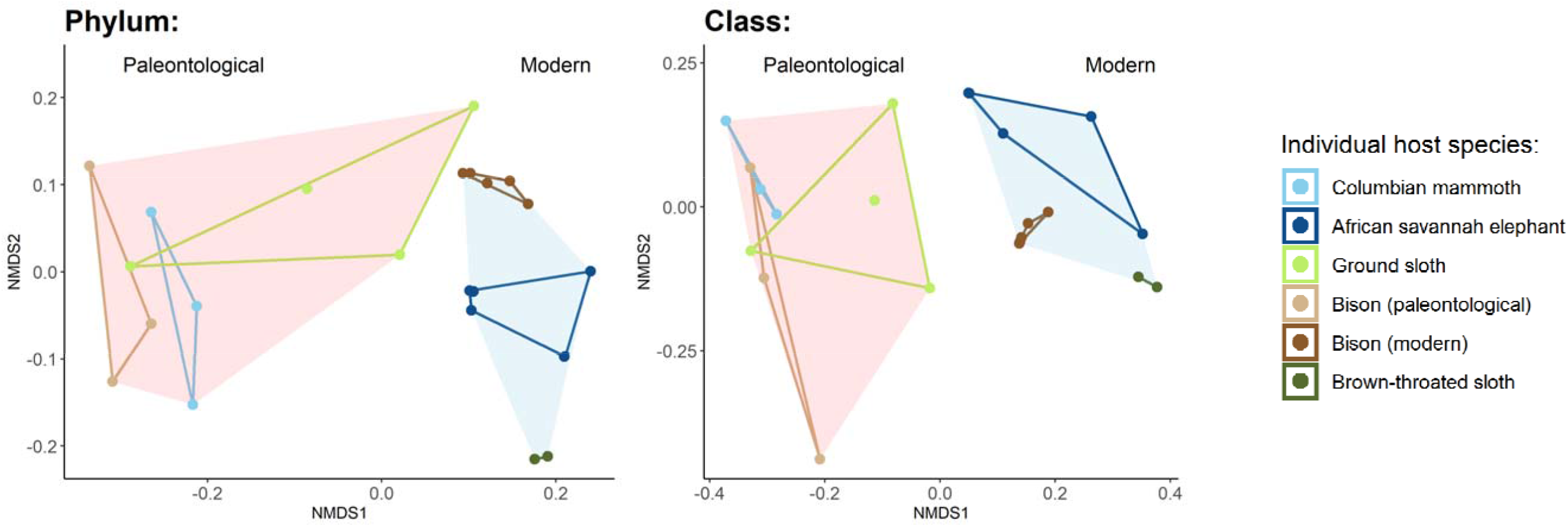
NMDS plots based on the Bray-Curtis distances between the bacterial communities in paleontological coprolites versus modern specimens at the phylum and class taxonomic levels. Samples from single host species are represented by distinctly colored points and are bound by blue and pink convex hulls, grouping the paleontological and modern samples, respectively.

### 4.5 BISON COPROLITES AND DESSICATION

NMDS based on the Bray-Curtis dissimilarities between the bacterial component of the paleontological and modern bison samples and cattle fecal matter from Wong et al (2016) at various stages of desiccation, showed that the cattle samples formed a continuum between modern and paleontological bison samples (Figure 4). Pairwise PERMANOVA between the bison and cattle samples at phylum level revealed that the bison coprolites were significantly different from cattle sample collected on days 0 – 8 (corrected p-values between 0.0246 and 0.0370 with individual p-values available in Supplementary Material), but from days 15 – 57 the cattle samples did not differ significantly from the paleontological samples (corrected p-values > 0.05). At class level, pairwise PERMANOVA showed that bison coprolites differed significantly from cattle samples collected on days 0 and 2 (corrected p-values 0.0261 and 0.0171, respectively) but from days 4 onwards did not differ significantly from the paleontological samples (corrected p-values > 0.05). Modern bison samples differed significantly from cattle samples on all days of desiccation post deposition and at both taxonomic levels but grouped most closely with cattle samples on days 2 – 6 at phylum and 2 – 4 at class (Figure 4).

**Figure 4.**
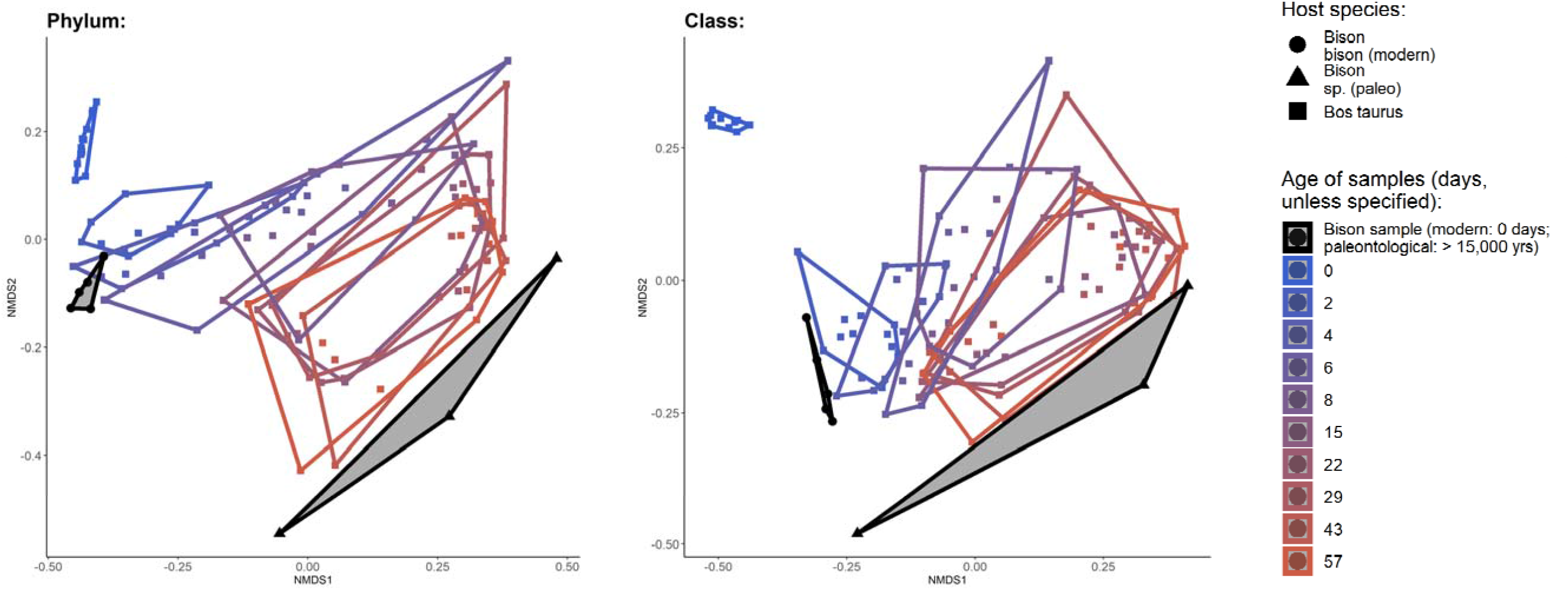
NMDS plots based on the Bray-Curtis distances between the bacterial component of *Bovidae* samples at the phylum and class taxonomic levels. Black points represent modern (circular), and paleontological (triangular) bison samples processed as part of this study, with each category bound by a convex hull with a semi-translucent fill. Colored points (square) are samples from Wong et al (2016), representing cattle fecal samples that were measured for bacterial composition at various days since deposition, from day 0 to 57, with each time symbolized by a different color and bound by a convex hull. Progressively desiccated cattle feces form a continuum between modern and paleontological bison samples when the diversity of the bacterial microbiome is assessed at phylum and class levels.

NMDS plots comparing bison samples processed in this study with the desiccated feces from Menke et al (2015) revealed a stark separation (Figure S3). While the springbok and giraffe samples formed a continuum from fresh to desiccated, these were distinct from the paleontological and modern bison classified here.

### 4.6 CONTAMINATION

To determine whether the coprolites were contaminated with significant levels of modern DNA, nucleotide transition rates were measured at the read termini using MapDamage2 (Jónsson et al. 2013) for bacterial phyla that had a reference genome. Only in paleontological bison samples was an increase in C-to-T transitions observed at the 5’ termini and corresponding A-to-G observed at the 3’ termini, compared to modern bison (Figure 5). The degradation and low concentration of endogenous DNA in paleontological samples, suggests that minimal modern contamination was introduced into the bison coprolites during laboratory processing.

**Figure 5.**
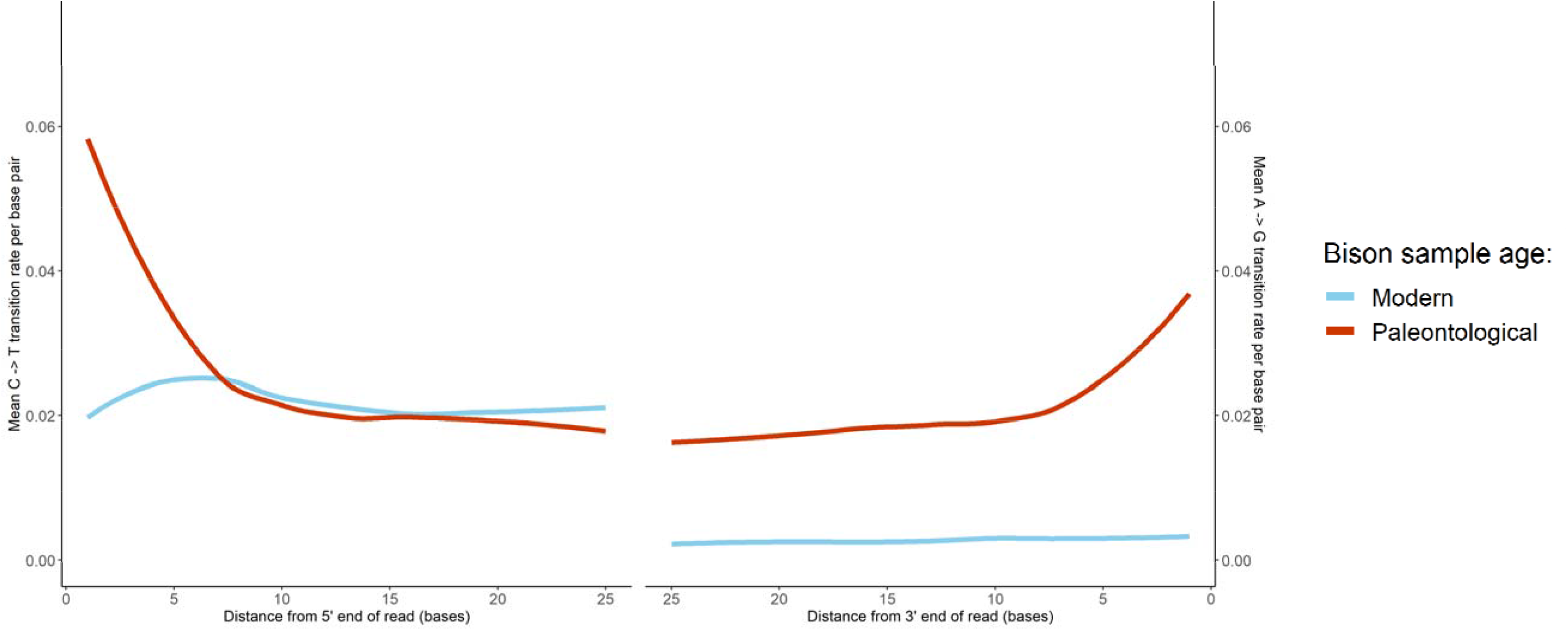
Rates of nucleotide transitions with distances form read termini for modern (blue) and palaeontologic (red) bison samples. The left-hand plot indicates the C-to-T transition rate with increasing distance from the 5’ read terminus, whereas the right-hand plot shows the A-to-G transition rate with increasing distance from the 3’ terminus.

## 5. DISCUSSION

Results from this study help answer a key question when evaluating the microbial composition of coprolites: – to what degree is it representative of the original gut microbiome versus the microbial changes that occur during desiccation? While the metagenomic readsets included in this study were separated by their paleontological versus modern classification, the convergence of progressively dried cattle feces (Wong et al. 2016) with bison coprolites indicates that this clustering may be in-part due to desiccation-related shifts in the community of gut microorganisms occurring in the weeks following coprolite deposition.

The convergence of microbiomes between desiccated cattle feces and bison coprolites was caused by common changes in the relative abundance (RA) of bacterial taxa. During the desiccation of cattle feces, Wong et al (2016) observed a decrease in the RA of Firmicutes and Bacteroidetes and an increase in Actinobacteria and Proteobacteria. A similar pattern was observed between the fresh and paleontological bison feces processed in this study. Fresh bison feces were dominated by Firmicutes, compared to the Actinobacteria that dominated the coprolites (Figure 2 and Table S3). At class level, bison coprolites and desiccated cattle feces both had a higher proportion of Actinobacteria, Alphaproteobacteria and Betaproteobacteria, and a lower proportion of Clostridia, compared to their fresh counterparts (Figure 2 and Supplementary Material).

A notable observation in the desiccation analysis was the separation of modern bison samples and fresh cattle feces. Modern bison samples were distinct from all the cattle specimens, but clustered most closely with cattle feces from days 2-6 (Figure 4). This difference could be due to a non-mutually exclusive combination of factors, including the domestication status (Metcalf et al. 2017), social structure (Grieneisen et al. 2017), contact patterns (Tung et al. 2015; VanderWaal et al. 2014; Blyton et al. 2014), diet (Groussin et al. 2017; Carmody et al. 2015; Degnan et al. 2012; Gordon and Cowling 2003; Kartzinel et al. 2019), or phylogenetic differences between hosts (Blekhman et al. 2015; Goodrich et al. 2014; Groussin et al. 2017), as well as the biases associated with comparing metagenomic and amplicon data (Durazzi et al. 2021). An improved desiccation study using metagenomic samples from bison feces collected on the Colorado Plateau and sampled according to Wong et al (2016) would reduce these confounding factors.

Another question that arises from a lack of desiccation research is whether variation between different species’ microbiomes in fresh fecal specimens is maintained through the drying process (Figure S4). If distinct communities of microorganisms at time zero tended to become more similar and loose beta diversity during desiccation, extrapolating the gut microbiome of extinct hosts from coprolites would be prevented (Figure S4: plot 3). However, microbe communities that maintained sufficient distinctiveness from one another despite desiccation and whose changes occur in a predictable direction, could be used to infer the original gut community of the host (Figure S4: plots 1, 2, and 4), and would open doors to information on extinct host traits through coprolite analysis.

The difference in results when comparing the bison samples processed here with samples from Wong et al (2016) versus Menke et al (2015) suggests that microbiomes from different species do not align during desiccation. The springbok and giraffe samples from Menke et al (2015) formed a continuum in the NMDS plots (Figure S3), from fresh to desiccated, but were distinct from the cattle, paleontological and modern bison samples. While springbok are also members of the Bovidae family, both species have different life-history traits, with springbok being mixed-feeders and giraffe obligate browsers. They also occupy different ecosystems, with different social groupings and interactions, all of which could influence microbiome composition. Nevertheless, Menke et al. (2015) dried samples for a shorter duration than Wong et al (2016) (7 versus 57 days, respectively), which may have resulted in springbok and giraffe microbiomes aligning with the bison and cattle samples given more time.

The current lack of fecal desiccation studies for species outside ungulates makes it difficult to determine whether the microbial community in modern elephant boluses would converge with mammoth coprolites. Large-scale, long-term, metagenomic desiccation studies across multiple modern species are needed and may allow researchers to determine whether factors such as gut type are detectable through time. If so, there would be potential to characterize traits from extinct species (without close relatives) that cannot be inferred from skeletal specimens, such as the digestive physiology of extinct ground sloths.

MTSv parameterized for the detection of aDNA may facilitate more studies assessing the gut microbiome of Late Pleistocene megafauna. MTSv is particularly well-suited for the classification of microbial species from paleontological samples as the assignment algorithm can align taxa that diverge from the genetic material within the reference database (*Tara Furstenau personal communication*; Furstenau et al. 2022). Previous methods for the rapid classification of metagenomic data, such as Kraken2 (D. E. Wood and Salzberg 2014), rely on exact k-mer matching. This can lead to low resolution of certain taxa and missed assignments when sequences diverge from the reference (*Tara Furstenau personal communication*; Furstenau et al. 2022).

Using MTSv to classify 200 firmicute species with a fragmentation and deamination pattern typical of paleontological samples indicated that USH are representative of taxon abundance at phylum, class, and order levels. At lower levels the USH counts do not reflect taxon abundances, preventing the subsequent use of abundance-based beta-diversity metrics and ordination analyses. Testing the relationship between MTSv USH and taxon abundance should be repeated for future desiccation studies of fresh fecal samples, as the correlation between USH and abundance is likely to weaken at lower taxonomic level, even in the absence of age-related fragmentation and deamination.

Radiocarbon dates proved the samples included in this study as some of the oldest published gut bacterial metagenomes analyzed from desiccated coprolites. The dates showed the bison and mammoth specimens as originating from the Late Pleistocene, with the Shasta ground sloth samples deposited in the Early Holocene. These results were consistent with radiocarbon dates from Mead and Agenboard (1992) for mammoth and Shasta ground sloth. However, the paleontological bison coprolites recorded here were older than Mead and Agenboard (1992) for samples collected at Grobot Grotto, and younger at Hooper’s Hollow. Only a few studies have assessed the gut microbiome of Late Pleistocene megafauna (> 11,700 ybp) using 16S rRNA amplicons. A notable study by Mardanov et al. (2012) evaluated the bacterial composition of the gut contents from a particularly well-preserved juvenile mammoth found in Siberia, named Lyuba (Kosintsev et al. 2010; Fisher et al. 2012). Those evaluating gut microbiomes from coprolites using metagenomic techniques have primarily focused on humans (Wibowo et al. 2021; Tito et al. 2012; Rivera-Perez et al. 2015; Cano et al. 2014; Rampelli et al. 2021a; Appelt et al. 2014) and dogs (Hagan et al. 2020; Borry et al. 2020) from the Late Holocene. A study by Bon et al (2012) used shotgun metagenomics to evaluate hyena coprolites that likely originated from the Late Pleistocene, but the authors focused on vertebrate rather than microbial DNA (Bon et al. 2012; Utge et al. 2020).

With studies of aDNA, the risk of contamination is considerable. Modern contamination introduced into the paleontological bison samples processed here was unlikely due to the increase in the C-to-T and A-to-G transitions detected at the 5’ and 3’ read termini, respectively. This pattern was not observed in reads from the coprolites of Shasta ground sloth, mammoth, or modern bison samples. These nucleotide changes were detected by extracting MTSv queries that mapped to a given taxon and aligning these against a reference genome for the same taxon downloaded from NCBI where available. At present, a limitation of this technique is the low percentage of bacterial species with reference genomes, but this limitation is likely to decrease as repositories continue to grow. The presence of older contamination is more difficult to assess as it would also have accumulated base pair transitions. However, desiccation of boluses from the arid environment would likely have limited microbial growth (Sinton et al. 2007; M. Chen and Alexander 1973; Yeager and Ward 1981; Rezaei and vander Gheynst 2010; Merino et al. 2019; Menke, Meier, and Sommer 2015; Wong et al. 2016). Over a two-year period, Walker et al. (2019) recorded a relative humidity of 50-55% at a dry cave from the Colorado Plateau (Walker et al. 2019), below the equilibrium relative humidity required for bacterial growth (60%) (Esbelin, Santos, and Hébraud 2018; Beuchat et al. 2013). The clustering of paleontological bison samples with desiccated cattle feces provided further evidence of minimal modern contamination. If present, genetic material from modern microbes would dominate the signal against the low concentration background of endogenous DNA.

The need to characterize paleontological microbiomes may increase as de-extinction efforts continue (DeFrancesco 2021). An understanding of mammoth gut microbiomes will be of importance in on-going efforts to reintroduce mammoth to northern latitudes, with the aim of re-establishing the mammoth-steppe grasslands. Coprophagy is well documented among young elephants to establish the cellulose digesting bacterial communities in the gut (Zimmer 2021). Similarly, juvenile mammoth specimens from permafrost have been found with GI tracts containing adult fecal material (Mardanov et al. 2012; Van Geel et al. 2011). At present, it is unclear whether boluses from extant Asian elephants would be effective in seeding the neo-mammoths with gut bacteria necessary for a life in the tundra.

## 6. CONCLUSION

This study categorized fecal metagenomes from Late Pleistocene megafauna using shotgun metagenomics and the novel MTSV classifier, showing that paleontological and modern fecal samples were clustered according to age and host species. The microbial composition of bison coprolites resembled modern desiccated cattle feces sampled at least 4 days post deposition. Future studies of coprolite metagenomes using shotgun metagenomics would benefit from running a parallel desiccation analysis of modern feces from a closely related species, wherever possible, to determine the degree to which the coprolite metagenome is a result of desiccation versus true dissimilarities between the modern and paleontological hosts. Also, a large-scale desiccation study including a variety of modern species may shed light on life-history traits of the extinct species without close extant relatives, by establishing the proximity of coprolite metagenomes and metagenomes of dried modern samples.

## 7. DISCLAIMER

Any opinions expressed in this paper are those of the authors and do not necessarily reflect the official positions and policies of the U.S. EPA and any mention of products or trade names does not constitute recommendation for use.

## Supporting information

Phyla present in samples by Kingdom

Classes present in samples by Phyla

Results of pairwise PERMANOVA between bison and cattle samples - taxonomic level: class

Results of pairwise PERMANOVA between bison and cattle samples - taxonomic level: phylum

