## Supplementary Material for "Metagenomic analysis of coprolites from three Late Pleistocene megaherbivores from the Southwestern United States"

### 1. SUPPLEMENTARY INFORMATION

#### COPROLITE COLLECTION SITES

**Bison sp**.: Mammoth Alcove is a medium-sized shelter at 1188 m elevation with a south-southeast exposure, whereas Grobot Grotto and Hooper's Hollow are both large south-facing alcoves close together, located at 1189 m and 1204 m elevation, respectively (Mead and Agenboard, 1989).

**Shasta ground sloth**: Two locations in Arizona, Muav and Rampart Caves, discovered in the early 1930s at an elevation of approximately 426 m and 530 m, and found to contain rich deposits of Late Pleistocene fossils and fecal material (Martin, Sabels and Shutler, 1961; Long and Martin, 1974; Hansen, 1978; Martin, Thompson and Long, 1985; Schmidt, Duszynski and Martin, 1992; Poinar *et al.*, 1998, 2003; Hofreiter *et al.*, 2000).

**Mammoth:** Bechan cave is a large sandstone grotto at 1280 m elevation, 52.8 m in depth, and identified in 1982 as containing a blanket layer of dry dung (Davis *et al.*, 1984; Mead *et al.*, 1986; Martin, 1987; Agenbroad and Mead, 1989; Mead and Agenboard, 1989).

### 2. SUPPLEMENTARY TABLES AND FIGURES

| **Species** | **Storage site** | **NPS* Accession number** | **Collection sites** | **Study identification numbers** | **Radiocarbon dates ± error** | **95% probability date ranges (ybp)** |
|---|---|---|---|---|---|---|
| Shasta ground sloth (*Nothrotheriops shastensis*) | Grand Canyon National Park Museum | 4597 | Rampart Cave, Mohave, AZ | GRCA-59574 | 10,927 ± 27 | 10,946 – 10,808 |
| | | | | | 11,240 ± 27 | 11,225 – 11,147 |
| | | | | | 10,873 ± 26 | 10,884 – 10,797 |
| | | | | | 11,351 ± 27 | 11,353 – 11,222 |
| | | | | | 11,338 ± 26 | 11,351 – 11,217 |
| | | 4712 | Muav cave, Mohave, AZ | GRCA-59511 | 11,180 ± 27 | 11,214 - 11,136 |
| | | | | | 11,446 ± 27 | 11,470 – 11,289 |
| | | | | GRCA-4712 | 11,232 ± 27 | 11,221 – 11,147 |
| | | | | | 11,412 ± 28 | 11,155 – 10,979 |
| | | | | | 11,412 ± 28 | 11,396 – 11,231 |
| | | | | | 11,093 ± 27 | 11,146 – 10,976 |
| | | | | | 11,231 ± 28 | 11,221 – 11,146 |
| Paleontological bison (*Bison sp.*) | Museum of Northern Arizona | 82 | Mammoth Alcove, Kane, UT | GLCA-821 | 18,611 ± 34 | 20,688 – 20,425 |
| | | | Grobot Grotto, Kane, UT | GLCA-877 | 18,806 ± 35 | 20,982 – 20,592 |
| | | | Hooper's Hollow, Kane, UT | GLCA-900 | 15,479 ± 30 | 16,903 – 16,772 |
| Columbian mammoth (*Mammuthus columbi*) | Museum of Northern Arizona | 81 | Bechan Cave, Kane, UT | GLCA-372 | 12,304 ± 29 | 12,856 – 12,143 |
| | | | | GLCA-370 | 12,456 ± 29 | 12,992 – 12,383 |
| | | | | GLCA-367 | 12,430 ± 30 | 12,923 – 12,352 |
| | | | | GLCA-2578 | 12,401 ± 30 | 12,896 – 12,315 |
| | | | | GLCA-382 | 12,426 ± 30 | 12,915 – 12,349 |
| | | | | GLCA-2627 | 12,427 ± 29 | 12,915 – 12,351 |

* NPS: National Park Service

**Table S1**. Summary of collection information, study identification numbers, AMS radiocarbon dates, as well as the number of reads and queries obtained from paleontological coprolite samples included in the study: Shasta ground sloth, paleontological bison, and Columbian mammoth (top to bottom). Multiple samples were run for Shasta ground sloth coprolites. The 95% date ranges were calculated using OxCAL version 4.4 (Ramsey, 2009) and atmospheric data from Reimer *et al.*, (2020).

| P1: Kmer query size | |
|---|---|
| 36 | 50 |

X

| P2: No edits | |
|---|---|
| 3 | 5 |

X

| P3: Binning mode | | |
|---|---|---|
| Fast | Efficient | Sensitive |

X

| P4: Minimum threshold for USH* | | | | | | | | | |
|---|---|---|---|---|---|---|---|---|---|
| 0 | 200 | 400 | 600 | 800 | 100 | 1200 | 1400 | 1600 | 1800 |

**Table S2**. The combination of parameters (P1-4) used for MTSv runs on 1,000,000 simulated gargammel aDNA reads. Kmer query length refers to the size of the deduplicated queries generated by the binning-analysis pipeline. Edits represents the threshold number of mismatches between a query and reference. Binning mode represents the combination of parameters (seed size, minimum seeds, and seed gap; parameter order respective hereafter) used by the assignment algorithm during binning. We used three binning modes during this study: fast (17, 5, 2), efficient (14, 4, 2), and sensitive (11, 3, 1). (* the USH threshold was converted into a proportion of the total USH generated by the MTSv run).

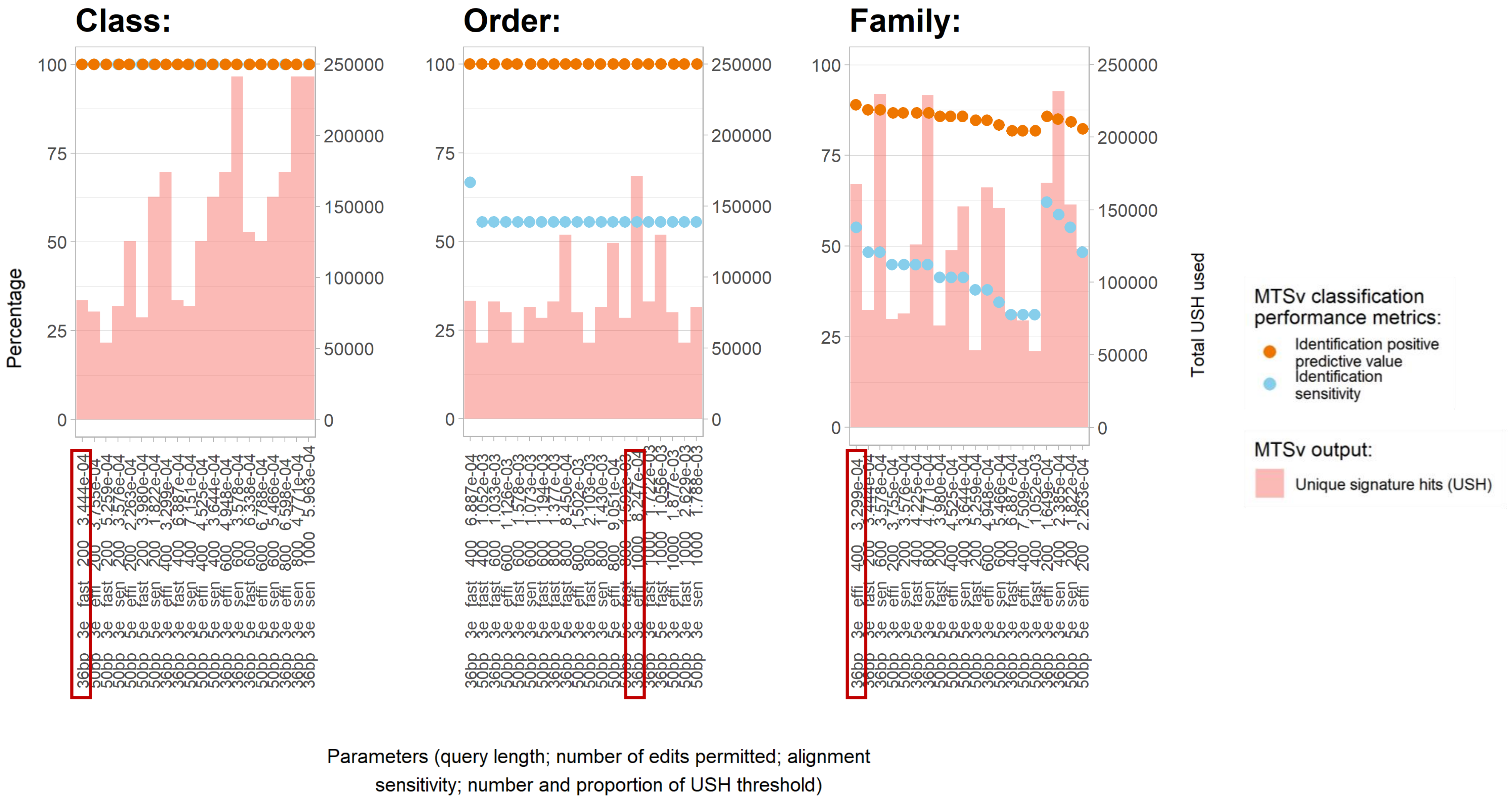

Class:
Order:
Family:
Percentage
Total USH used
MTSv classification performance metrics:
Identification positive predictive value
Identification sensitivity
MTSv output:
Unique signature hits (USH)
Parameters (query length; number of edits permitted; alignment sensitivity; number and proportion of USH threshold)

**Figure S1**. The top 20 MTSv runs in identifying the classes, orders and families associated with the reads from 200 firmicute species processed by gargammel to include age-related changes. These runs were ranked in descending order of positive predictive values (orange points) and sensitivity scores (blue points). MTSv runs differed in the query lengths, number of edits permitted between a query and reference, the sensitivity of alignment (determined by alignment parameters summarized in Table S2), and the minimum number of USH for taxon inclusion (converted to a proportion of the total USH), as well as the total number of USH used by the classifier (pale pink histogram). Highlighted MTSv runs indicate the parameter combination selected for further downstream analyses of coprolites (36bp query length, 3 edits between query and reference and efficient alignment, with USH thresholds varying for different taxonomic levels).

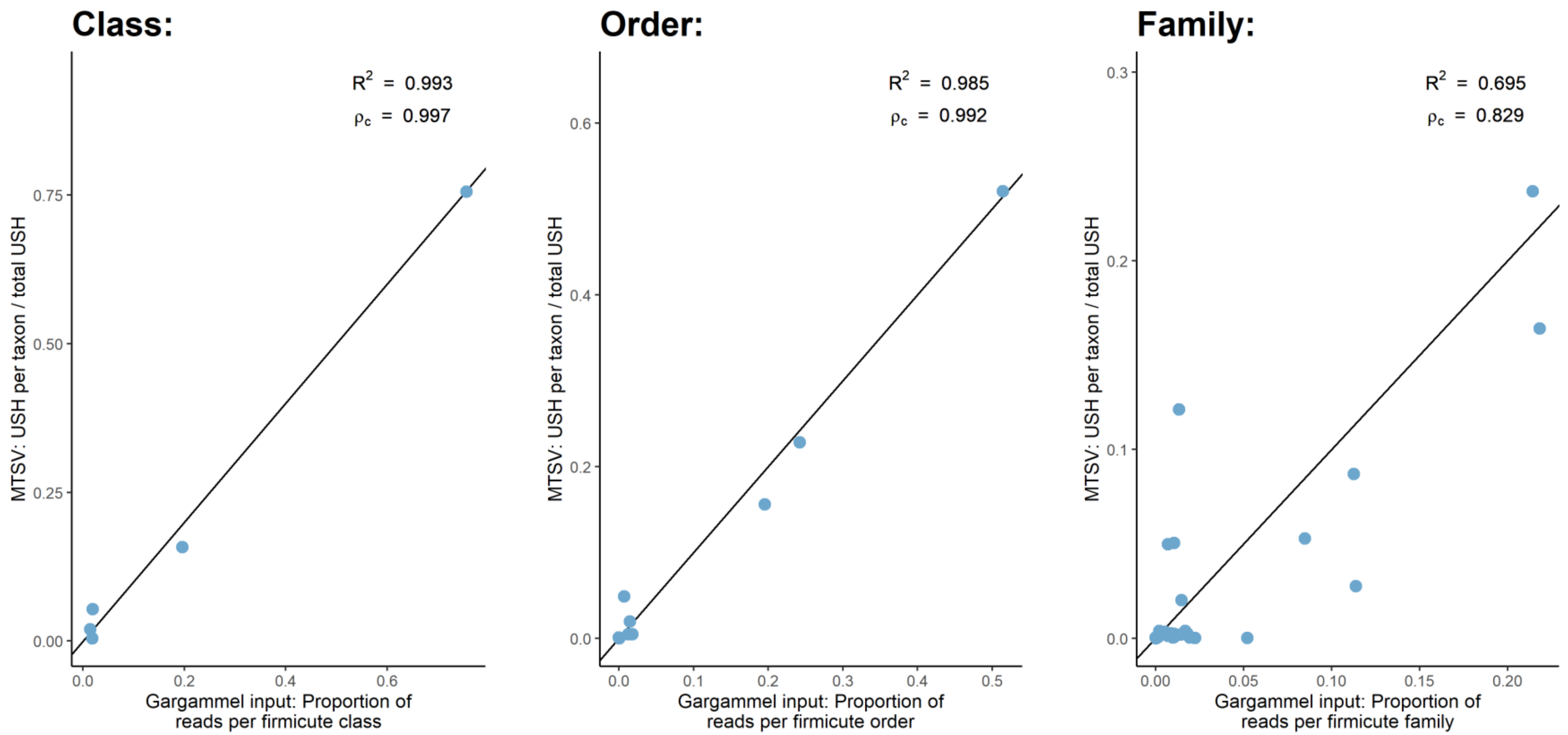


**Figure S2**. The relative abundance of gargammel reads from each taxon versus the proportion of USH identified by MTSv at the class, order, and family levels. The points displayed came from the MTSv run parameterized for a 36 bp query length, 3 edits and an efficient alignment mode. Each comparison shows the $R^2$ value of the points away from a 1-to-1 line as well as Lin's concordance correlation coefficient ($\rho_c$). The figure highlights that MTSv USHs are a good indicator of bacterial abundance at taxonomic levels above family.

| Species | Kingdom | Phylum: percentage |
|---|---|---|
| Bison *bison* (modern) | Archaea | Euryarchaeota: 13.1743890722925 |
| | Bacteria | Firmicutes: 67.5520625769282,<br>Bacteroidetes: 7.44743119898517,<br>Actinobacteria: 4.20642352093039,<br>Proteobacteria: 2.64228434768255 |
| | Eukaryota | Streptophyta: 2.98733840850779 |
| Bradypus *variegatus* | Archaea | Euryarchaeota: 7.82385380631297 |
| | Bacteria | Verrucomicrobia: 35.3748023500362,<br>Bacteroidetes: 34.9505239153169,<br>Firmicutes: 9.5333940320368,<br>Proteobacteria: 8.00912507747584,<br>Lentisphaerae: 2.60800869647104,<br>Actinobacteria: 1.09013993949783 |
| Loxodonta *africana* | Archaea | Euryarchaeota: 1.53448308845536 |
| | Bacteria | Proteobacteria: 47.1217352111144,<br>Bacteroidetes: 16.5815399968847,<br>Firmicutes: 16.1444151957561,<br>Verrucomicrobia: 3.49460616137364,<br>Actinobacteria: 3.46800601395302,<br>Spirochaetes: 1.36889802527466,<br>Tenericutes: 1.02575840554082 |
| | Eukaryota | Streptophyta: 5.96165209225422 |
| Bison *sp.* (paleontological) | Bacteria | Actinobacteria: 49.9242291884496,<br>Proteobacteria: 34.2753071984631,<br>Firmicutes: 15.4635984544028 |
| Mammuthus *columbi* | Bacteria | Actinobacteria: 51.66045226374,<br>Proteobacteria: 44.5132985506531,<br>Firmicutes: 3.47683619907833 |
| Nothrotheriops *shastensis* | Bacteria | Actinobacteria: 39.8043818948311,<br>Firmicutes: 37.8428912509215,<br>Proteobacteria: 14.7871067207959,<br>Bacteroidetes: 6.25992283764298 |

**Supplementary Table 3**. The percentages of the dominant phyla (>1% USH) per kingdom for each paleontological and modern species. The majority of USHs from paleontological samples were assigned to Actinobacteria and Proteobacteria, with a high percentage of Firmicutes detected in the coprolites of N. *shastensis*.

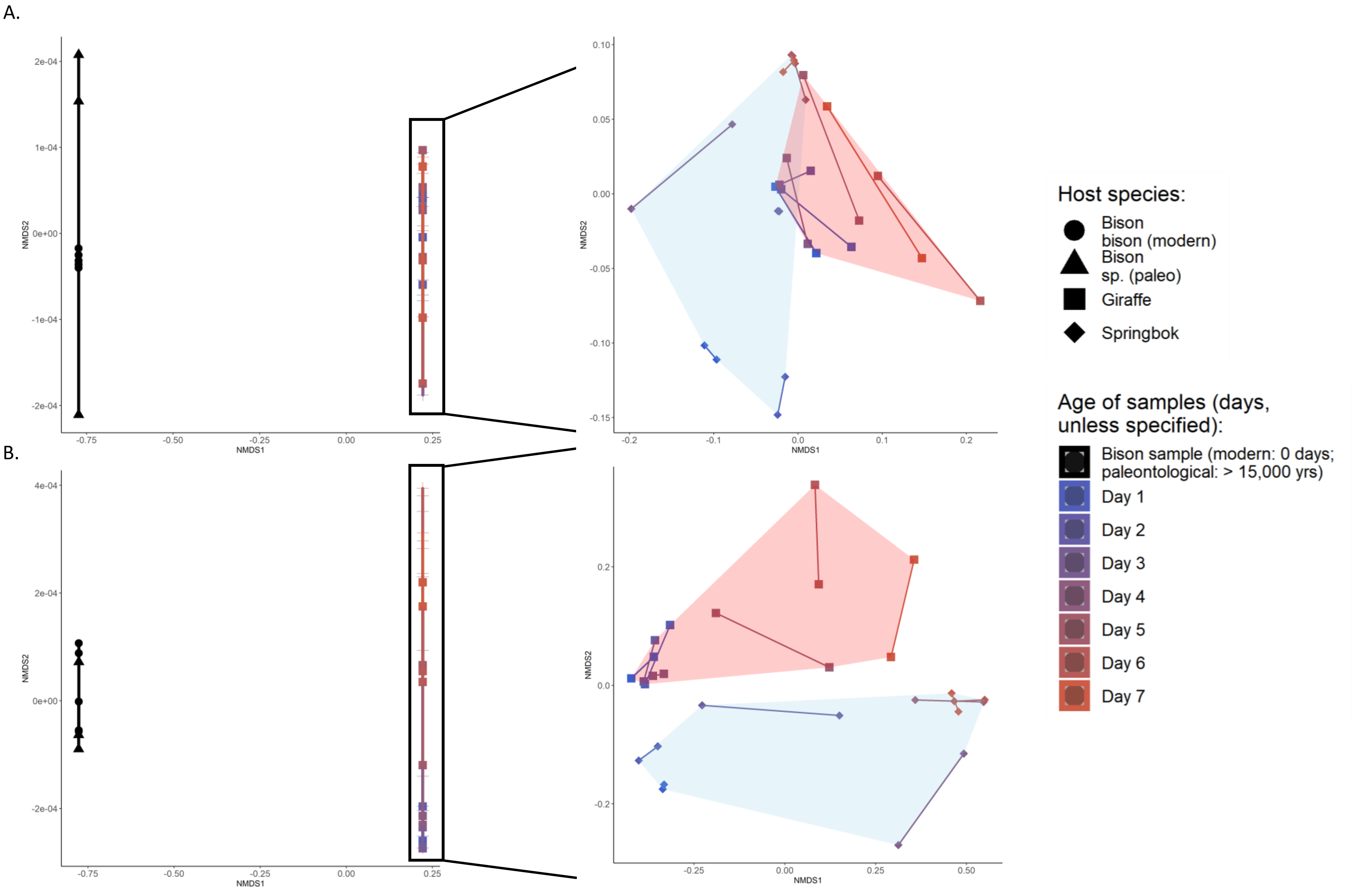
A.
B.
NMDS1
NMDS2
Host species:
Bison bison (modern)
Bison sp. (paleo)
Giraffe
Springbok
Age of samples (days, unless specified):
Bison sample (modern: 0 days; paleontological: > 15,000 yrs)
Day 1
Day 2
Day 3
Day 4
Day 5
Day 6
Day 7

**Figure S3**. The left-hand NMDS plots show the dissimilarity between paleontological bison, modern bison, and desiccated samples from Menke et al (2015) at the phylum (A) and class (B) levels. For each taxonomic level, the right-hand NMDS plots show the Menke et al (2015) giraffe and springbok samples (pink and blue convex hulls, respectively) after ordination was repeated without the bison samples. This was done to show the separation of points along a continuum from fresh to desiccated.

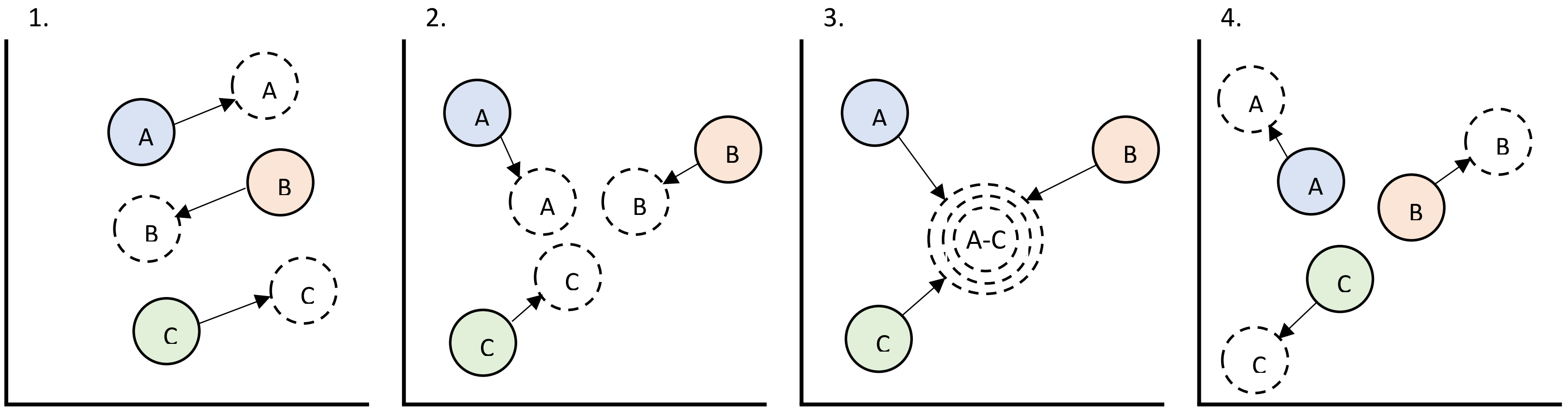


**Figure S4**. Hypothetical ordination plots showing how the distinctiveness of microbiomes in fresh fecal samples from different host species (clusters represented as solid shaded circles A - C) might change during desiccation studies (clusters as dashed circles A – C). The four plots represent situations during desiccation in which the beta diversity of the three samples: 1) does not change, 2) converges somewhat, 3) converges completely, and 4) increases. In plots 1, 2, and 4, the original microbiome may be inferred from the desiccated samples. However, the complete convergence of desiccated microbiomes in plot 3 would prevent this.
